# E-cadherin binds glycosylated sodium-taurocholate cotransporting polypeptide to facilitate hepatitis B virus entry

**DOI:** 10.1101/729822

**Authors:** Qin Hu, Fei-Fei Zhang, Liang Duan, Bo Wang, Pu Li, Dan-Dan Li, Sheng-Jun Yang, Lan Zhou, Wei-Xian Chen

## Abstract

Hepatitis B virus (HBV) continues to pose a serious public health risk and is one of the major causes of chronic liver disease and hepatocellular carcinoma. Current antiviral therapy does not effectively eradicate HBV and, thus, further investigation into the mechanisms employed by HBV to allow for invasion of host cells, is critical for the development of novel therapeutic agents. Sodium-taurocholate cotransporting polypeptide (NTCP) has been identified as a functional receptor for HBV. However, the specific mechanism by which HBV and NTCP interact remains unclear. Herein we show that the expression of E-cadherin was upregulated in cells expressing HBV, while knockdown of E-cadherin in HepG2-NTCP cells, HepaRG cells and primary human hepatocytes served to significantly inhibit infection by HBV and HBV pseudotyped particles. Alternatively, exogenous E-cadherin expression was found to significantly enhance HBV uptake by HepaRG cells. Further, mechanistic studies identified glycosylated NTCP localized to the cell membrane via E-cadherin binding, which subsequently allowed for more efficient binding between NTCP and the preS1 of the large HBV surface proteins. E-cadherin was also found to play a key role in establishing and maintaining hepatocyte polarity, which is essential for efficient HBV infection. These observations suggest that E-cadherin facilitates HBV entry through regulation of NTCP distribution and hepatocyte polarity.

**Author Summary:** Hepatitis B Virus (HBV) still seriously endangers public health. It is very important to understand the mechanism of HBV invading host cells for developing new therapy target. Sodium-taurocholate cotransporting polypeptide (NTCP) is the key receptor mediating HBV invasion, while other molecules also exhibit important roles in ensuring efficient and productive HBV infection. This study reports that E-cadherin facilitates HBV entry by directly interacting with glycosylated NTCP to mediate its distribution on the hepatocyte membrane and also affects the efficacy of HBV invasion by influncing hepatocyte polarity.

## Introduction

Hepatitis B virus (HBV), a member of the *Hepadnaviridae* family, is a small, enveloped DNA virus [1] that infects human liver parenchymal cells. Although the widespread use of vaccines has greatly reduced the rate of infection, HBV continues to pose a serious threat to global health, affecting more than 350 million individuals worldwide, all of whom are at increased risk of developing liver cirrhosis and hepatocellular carcinoma (HCC) [2]. Current therapeutic regimens that employ direct-acting antivirals, with or without ribavirin, have significantly increased the prevalence of escape mutants and cause serious adverse effects, thus, eliciting a low curative rate in HBV patients [3].

Furthermore, the lack of effective therapeutic options for HBV is partially due to our incomplete understanding of the HBV life cycle, including the stages during which the virus enters host cells, undergoes DNA replication, and assembles and releases virions from the cells. Productive HBV infection occurs following viral entry into a host cell, which is initiated through the binding of the preS1 domain of the large HBV surface proteins (LHBs) with high affinity HBV receptors on hepatocytes [4,5]. Recently, the sodium-taurocholate cotransporting polypeptide (NTCP), a bile acid transporter expressed at the basolateral membrane of human hepatocytes, has been identified as a functional receptor for HBV [6]. Exogenous expression of NTCP in human hepatoma cell lines, such as HepG2 or Huh7, confers susceptibility to HBV infection, and thus, constitute effective cell culture models for examining HBV entry [6,7]. However, overexpression of NTCP in human extrahepatic cell lines, such as HeLa cells, or in mouse hepatocyte cell lines, such as Hepa1-6 and MMHD3 cells, is not enough to allow for competent HIV infection of these cells, suggesting that additional molecules are also required for efficient HBV infection [8].

An additional study revealed that overexpression of the hepatitis B surface antigen binding (SBP) protein in HepG2 cells (HepG2-SBP) induced susceptibility to HBV infection. SBP was shown to interact directly with HBV envelop proteins [9]. Moreover, heparin sulfate proteoglycans (HSPGs) have been described as attachment receptors for HBV at the surface of hepatocytes and function to bring the virus into close proximity with NTCP [10]. Verrier et al. also reported that Glypican 5 attaches to the surface of HBV particles prior to NTCP binding, thereby assisting in host cell entry [11]. These studies indicated that, in addition to NTCP, other molecules exhibit important roles in ensuring efficient and productive HBV infection.

Cadherin adhesion molecules are core components in adherens junctions, located on the basement membrane aspect of polarized epithelial cells. E-cadherin, specifically, is a calcium-dependent adhesion integrin that is abundant in epithelial tissues and plays an important role in cell–cell adhesion complexes including desmosomes and adherens junctions [12]. In our study, we found significantly increased expression of E-cadherin in cells that stably or transiently expressed HBV, which is inconsistent with studies that have reported that hepatitis B virus X protein (HBx) represses E-cadherin expression [13]. These findings suggest that E-cadherin may have an undefined role in the HBV life cycle. Furthermore, cell adhesion molecules are highly associated with viral invasiveness and, thus, we hypothesized that E-cadherin may be associated with efficient HBV entry. Herein, we investigated the precise role, and subsequent mechanism, employed by E-cadherin in modulating HBV infection.

## Results

### E-cadherin is upregulated in HBV-expressing cells

To identify the effect of HBV replication on E-cadherin, we firstly examined the expression of E-cadherin in HepG2, HepG2.2.15 (stably express HBV), and HepG2 cell lines that had been transiently transfected with the HBV-expressing plasmid pcDNA3.1-HBV1.1 or pcDNA3.1-HBV1.3 or vector pcDNA3.1. The results showed that HBV expression and replication promoted expression of E-cadherin (Fig 1A and 1C). We also examined changes in E-cadherin expression in HepAD38 cells treated with tetracycline, and HepG2.2.15 cells transfected with siRNA targeting HBV. Results show that E-cadherin was downregulated following inhibition of HBV replication (Fig 1B and 1C). These results suggests that E-cadherin was significantly upregulated by HBV expression, suggesting that E-cadherin may contribute to the lifecycle of HBV.

**Fig 1.**
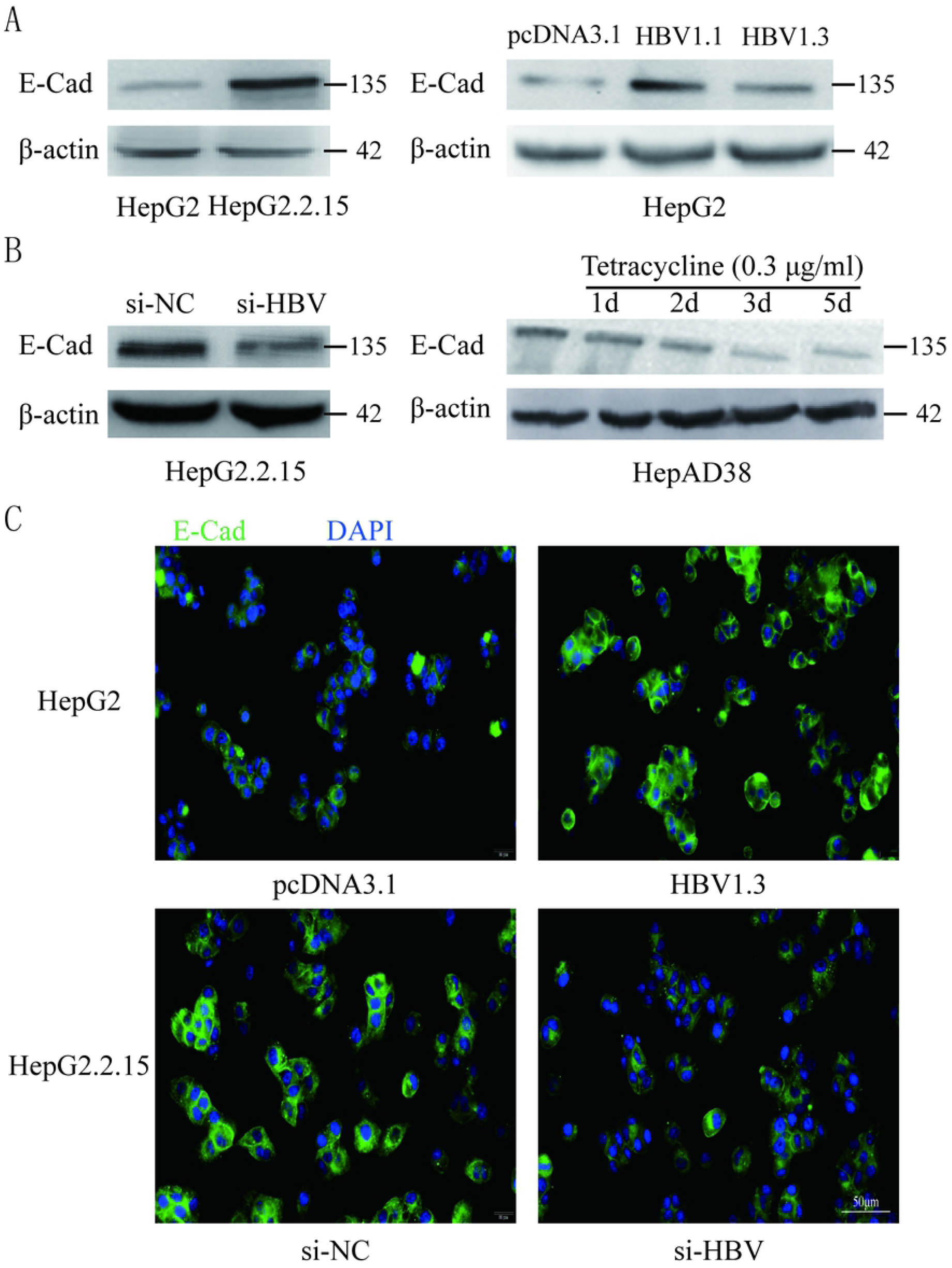
The expression of E-cadherin in cells that stably or transiently express HBV. (A) Expression of E-cadherin in HepG2 and HepG2.2.15 cells (left panel) or HepG2 cells that had been transiently transfected with the pcDNA3.1-HBV1.1, pcDNA3.1-HBV1.3 or pcDNA3.1 plasmids (right panel) were analyzed via western blot analysis. β-actin was used as a loading control. (B) Expression of E-cadherin in HepG2.2.15 cells transfected with siRNA-NC or siRNA-HBV (left panel) or HepAD38 cells cultured in media containing 0 μg/mL or 0.3 μg/mL of tetracycline (right panel). (C) E-cadherin expression as determined by immunofluorescence in HepG2 cells transiently transfected with vector pcDNA3.1 (pcDNA3.1) or plasmid pcDNA3.1-HBV1.3 (HBV1.3); and HepG2.2.15 cells transfected with siRNA-NC (si-NC) or siRNA-HBV (si-HBV). Representative data was shown from triplicate experiments. E-Cad: E-cadherin.

### Downregulation of E-cadherin reduces infection efficacy of HBV particles

To better elucidate the role that E-cadherin has in modulating HBV infection, we performed various virologic assays. HepG2-NTCP, HepaRG, and PHH cells were transfected with siRNA-NC, siRNA-E-cadherin, siRNA-NTCP or siRNA-E-cadherin together with siRNA-NTCP before being infected with HBV particles, which had previously been enriched from the supernatant of HepAD38 cells. The mRNA level of E-cadherin and NTCP were significantly downregulated following siRNA treatment in HepG2-NTCP and HepaRG cell lines (Fig 2A and 2B). Moreover, silencing of E-cadherin and NTCP significantly reduced the level of HBV 3.5 kb mRNA in HepG2-NTCP, HepaRG and PHH cell lines (Fig 2C); while silencing of both E-cadherin and NTCP in HepG2-NTCP and PHH cells served to further reduce the level of HBV 3.5 kb mRNA compared to that observed by either E-cadherin or NTCP separately. Further, western blot and immunofluorescence analysis determined that downregulation of E-cadherin or NTCP independently, or together, significantly reduced the expression of HBV core protein and HBV HBsAg with similar inhibition efficiencies (Fig 2D and 2E). These results suggest that silencing of E-cadherin significantly inhibited the infection of HBV particles enriched from the supernatant of HepAD38 cells.

**Fig 2.**
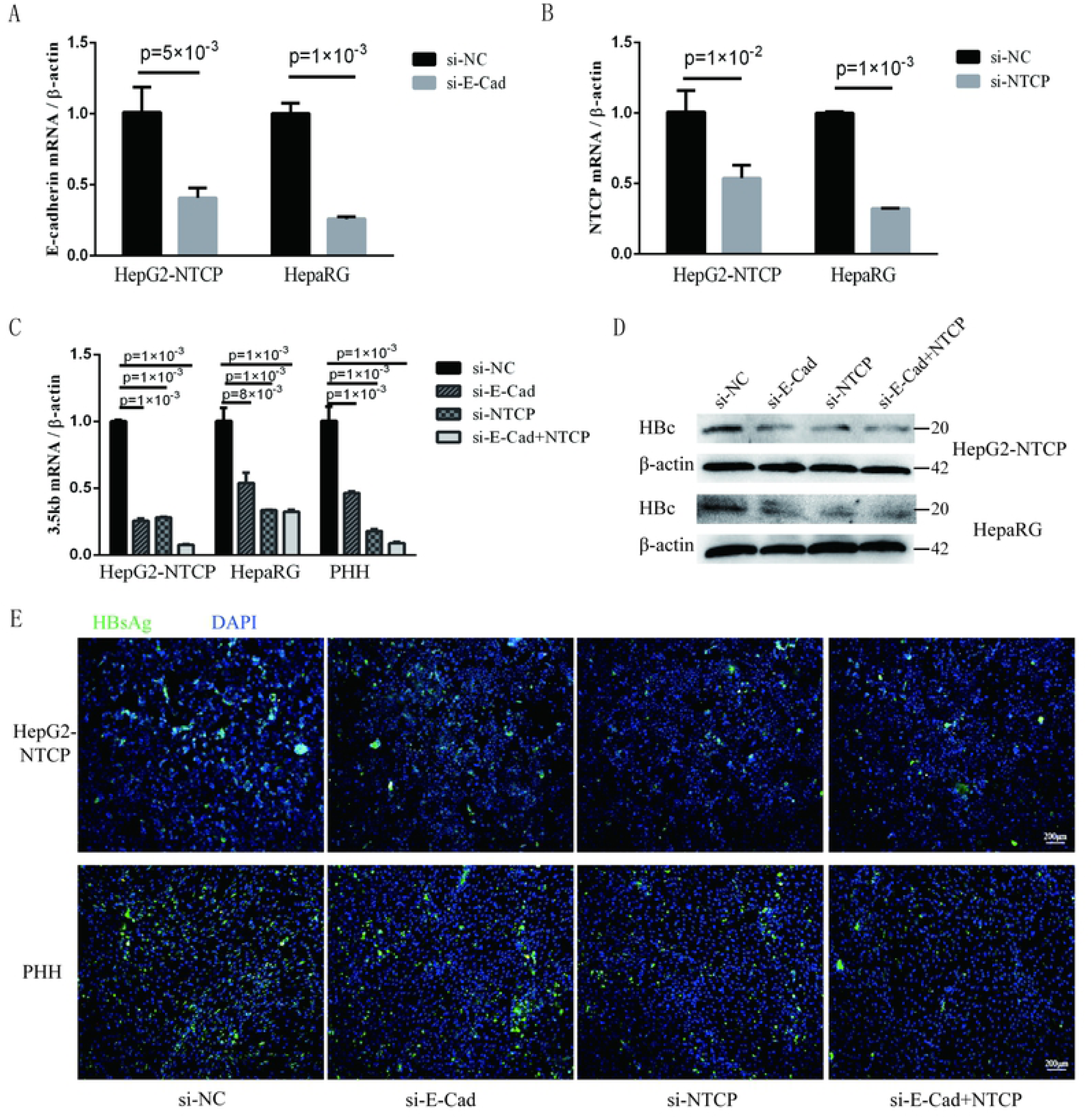
Downregulation of E-cadherin reduces infection efficacy of HBV particles enriched from supernatants of HepAD38 cells. (A) E-cadherin expression was silenced in HepG2-NTCP and HepaRG cells by siRNAs. (B) NTCP expression were silenced in HepG2-NTCP and HepaRG cells by siRNAs. Silencing efficacy was assessed for E-cadherin and NTCP by qRT-PCR two days post-transfection. (C) Total RNA was extracted two days post-infection and HBV infection was assessed by qRT-PCR quantification of HBV 3.5 kb mRNA normalized to β-actin mRNA. (D) Total protein was extracted three days post-infection and expression of HBV core (HBc) was assessed by western blot analysis in HepG2-NTCP and HepaRG cells. (E). HBsAg expression was assessed by immunofluorescence three days post-infection in Hep2-NTCP and PHH cells treated with si-NC, si-E-cadherin, si-NTCP or si-E-cadherin and si-NTCP together. Representative data is shown from triplicate experiments.

### Silencing of E-cadherin inhibits infection with HBV particles isolated from the serum of an HBV carrier

To further elucidate the relationship between E-cadherin and HBV infection, HepG2-NTCP and PHH cells were infected with HBV particles obtained from a chronic HBV patient. Infection were detected after four days. Silencing of E-cadherin or NTCP alone or at the same time, acted to significantly reduce the level of HBV 3.5 kb mRNA in HepG2-NTCP and PHH cells (Fig 3A). Moreover, silencing of E-cadherin also significantly inhibited the level of HBV core and HBsAg proteins (Fig 3B and 3C). These results suggest that silencing of E-cadherin also inhibits infection by HBV particles isolated from the serum of HBV carriers.

**Fig 3.**
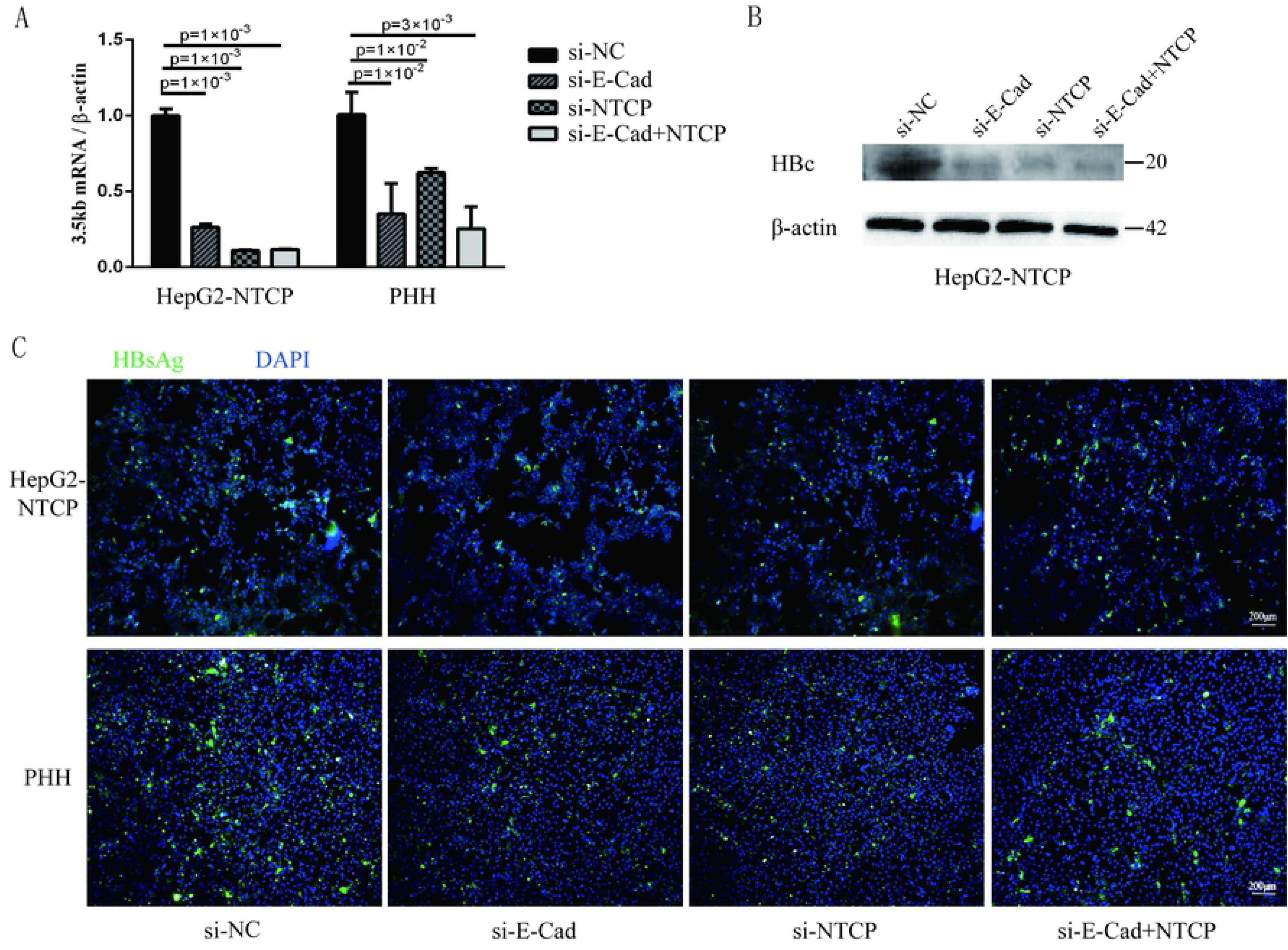
Downregulation of E-cadherin inhibits infection of HBV particles from the serum of an HBV carrier. HepG2-NTCP and PHH cells were incubated with the HBV-positive serum diluted in DMEM at a MOI of 10 after three days post-transfection with si-NC, si-E-cadherin, si-NTCP or si-E-cadherin and si-NTCP together. (A) Total RNA was extracted four days post-infection and HBV infection was assessed via qRT-PCR quantification of HBV 3.5 kb mRNA normalized to β-actin mRNA. (B) Total protein was extracted four days post-infection and HBV core (HBc) expression was assessed by western blot analysis in HepG2-NTCP cells. (C) HBsAg expression was assessed by immunofluorescence four days post-infection in Hep2-NTCP and PHH cells. Representative data is shown from triplicate experiments.

### Overexpression of E-cadherin promotes HBV particles infection

The concentration of E-cadherin was determined to be lower in HepaRG cells compared to HepG2-NTCP cells. Therefore, to elucidate the effect of E-cadherin overexpression on HBV infection, HepaRG cells were transfected with either pcDNA3.1-E-cadherin (E-cadherin) or pcDNA3.1 (vector). The transfection efficiency of E-cadherin was confirmed via western blot analysis (Fig 4A). At three days post-transfection, HepaRG cells were infected with enriched HBV particles. E-cadherin overexpression increased the level of HBV 3.5 kb mRNA (Fig 4B). Furthermore, overexpression of E-cadherin served to enhance the protein level of HBV core and HBsAg proteins (Fig 4C and 4D). These results suggest that E-cadherin contributed to HBV infection.

**Fig 4.**
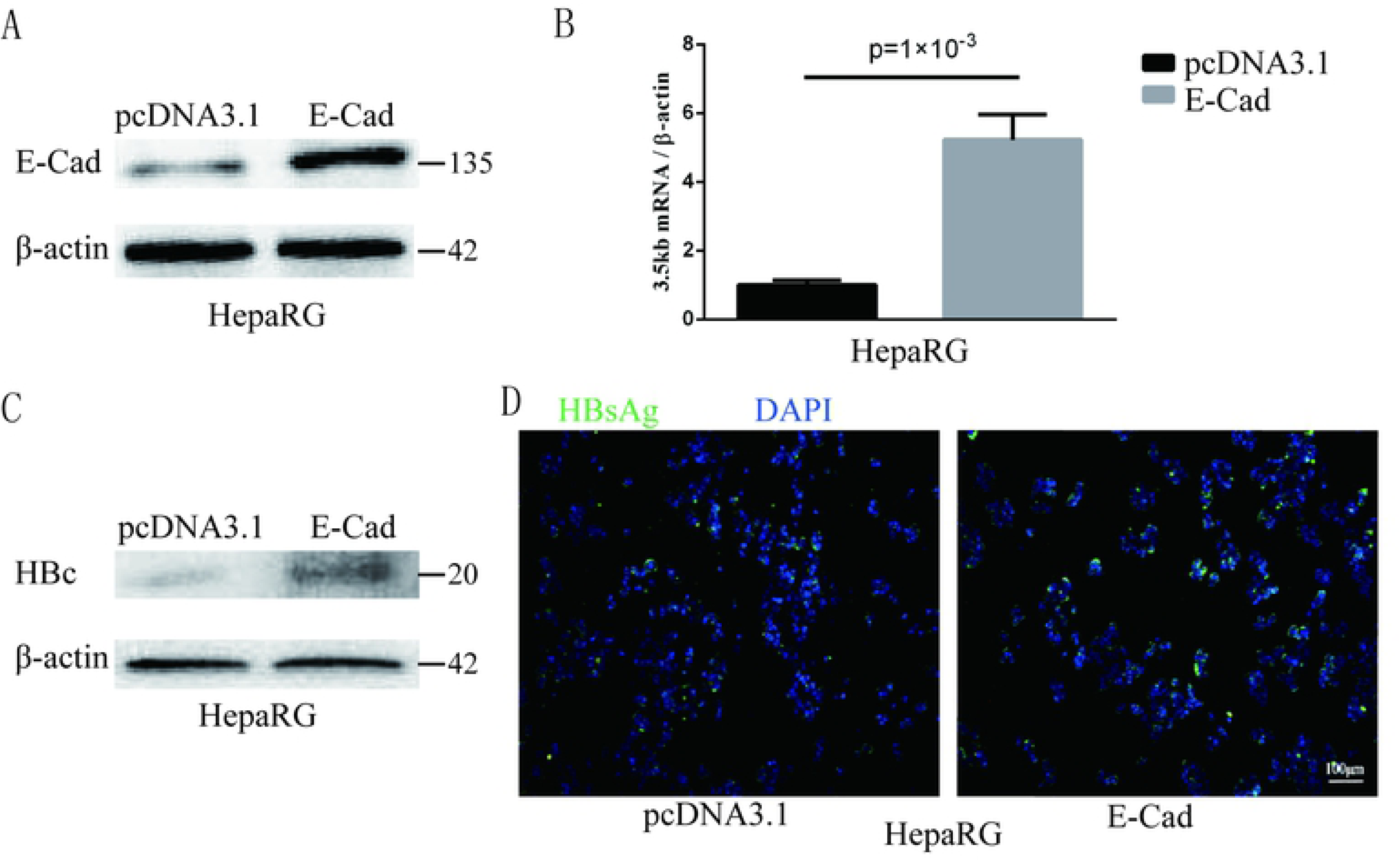
Overexpression of E-cadherin promotes HBV particles infection in HepaRG cells. (A) E-cadherin expression was assessed via western blot analysis three days post-transfection with pcDNA3.1 or pcDNA3.1-E-cadherin plasmids in HepaRG cells. (B) Total RNA was extracted two days post-infection and the amount of HBV was assessed via qRT-PCR quantification of HBV 3.5 kb mRNA normalized to β-actin mRNA. (C) Total protein was extracted three days post-infection and HBV core (HBc) expression was assessed by western blot analysis in HepaRG cells. (D) HBsAg expression was assessed by immunofluorescence three days post-infection in HepaRG cells. Representative data is shown from triplicate experiments. E-Cad: E-cadherin.

### E-cadherin specifically modulates HBV pseudoparticles entry

To further clarify the steps of the HBV life cycle impacted by E-cadherin expression, viral entry assays were conducted using HBVpps, a pseudotyped virus that expresses the HBV large protein. Silencing of E-cadherin or NTCP individually or together by siRNAs served to significantly inhibit HBVpps entry into HepG2-NTCP and HepaRG cells (Fig 5A and 5B), suggesting that E-cadherin impacts HBV binding/entry rather than HBV replication.

**Fig 5.**
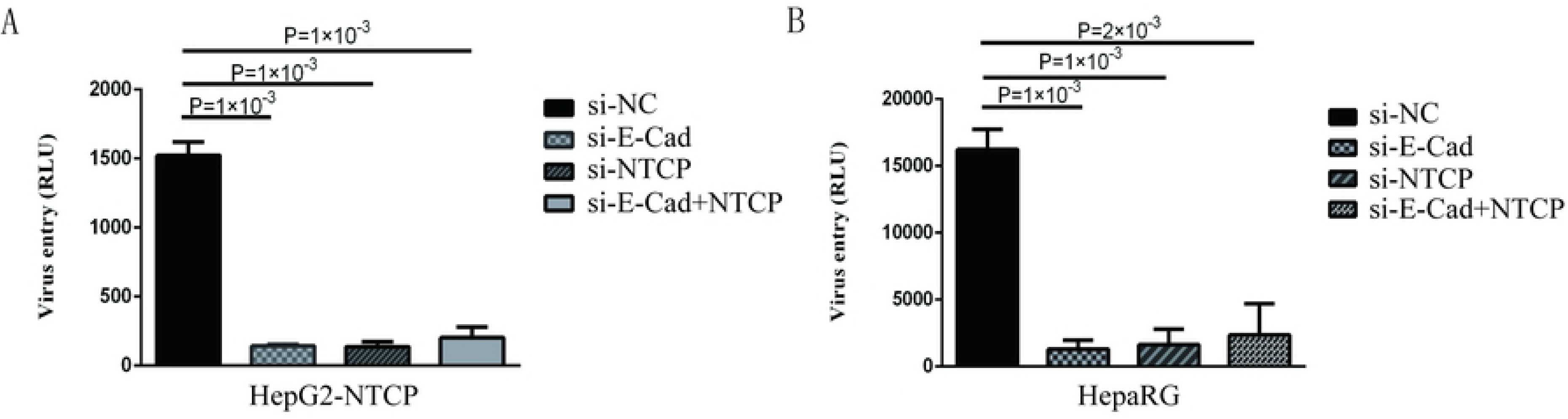
Downregulation of E-cadherin inhibits entry of HBV pseudoparticles. (A, B)HepG2-NTCP and HepaRG cells were infected with luciferase-encoding pseudotyped virus particles bearing HBV large envelope glycoprotein LHBs three days post-transfection with siRNA. Luciferase assays were performed 72 h after infection. Representative data is shown from triplicate experiments.

### Silencing of E-cadherin inhibits HBV pre-S1 binding and internalization to HepG2-NTCP and PHH cells

We next sought to determine the mechanism employed by E-cadherin to enhance HBV binding. We, therefore, quantified the expression of HBV pre-S1 (Myr-2-47aa) via immunofluorescence assays in HepG2-NTCP and PHH cells after 120 min incubation at 4 °C, which is the conditions in which viral binding most readily occurs, or at 37 °C, when uptake of pre-S1 occurs. Results revealed that when E-cadherin or NTCP were silenced separately or at the same time, significant inhibition of preS1 binding and uptake were observed in HepG2-NTCP and PHH cells (Fig 6A and 6B). These results suggest that E-cadherin modulates HBV entry by affecting preS1 binding and internalization by host cells.

**Fig 6.**
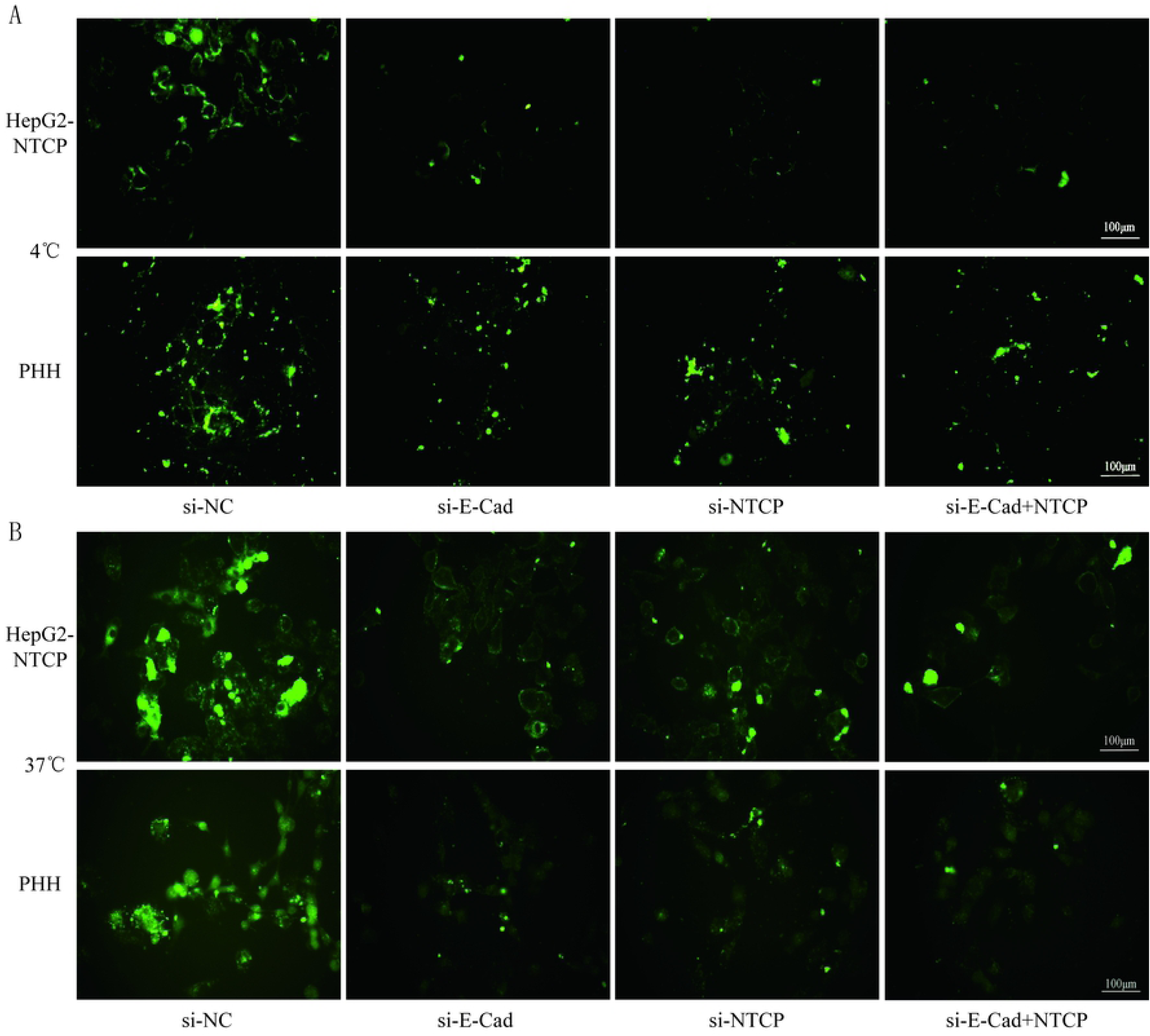
Silencing E-cadherin inhibits HBV preS1 binding and internalization by HepG2-NTCP and PHH cells. HepG2-NTCP and PHH cells were cultured with HBV preS1 for 2 h at (A) 4 °C or (B) 37 °C three days after transfected with si-NC, si-E-cadherin, si-NTCP or si-E-cadherin and si-NTCP together. Representative data is shown from triplicate experiments.

### E-cadherin regulates NTCP cell surface distribution

To further explore the mechanism by which E-cadherin mediates HBV particle entry, we examined whether it influenced the total intracellular NTCP concentration. Results show that silencing of E-cadherin did not affect intracellular NTCP in HepG2-NTCP and HepaRG cells (Fig 7A and 7B), which suggests that E-cadherin does not modulate HBV entry by directly affecting the expression and stability of NTCP. We, therefore, speculated that E-cadherin may influence the membrane distribution of NTCP. Our results clearly shown that silencing E-cadherin in HepG2-NTCP cells significantly impacted the subcellular distribution of NTCP. Specifically, NTCP which, primarily localizes to the cell surface was instead transferred to the cytoplasm (Fig 7C). We next separated membrane proteins to confirm the observed changes in NTCP cell surface distribution. As shown in Fig 7D, knockdown of E-cadherin resulted in significantly reduced levels of NTCP at the cell membrane.

**Fig 7.**
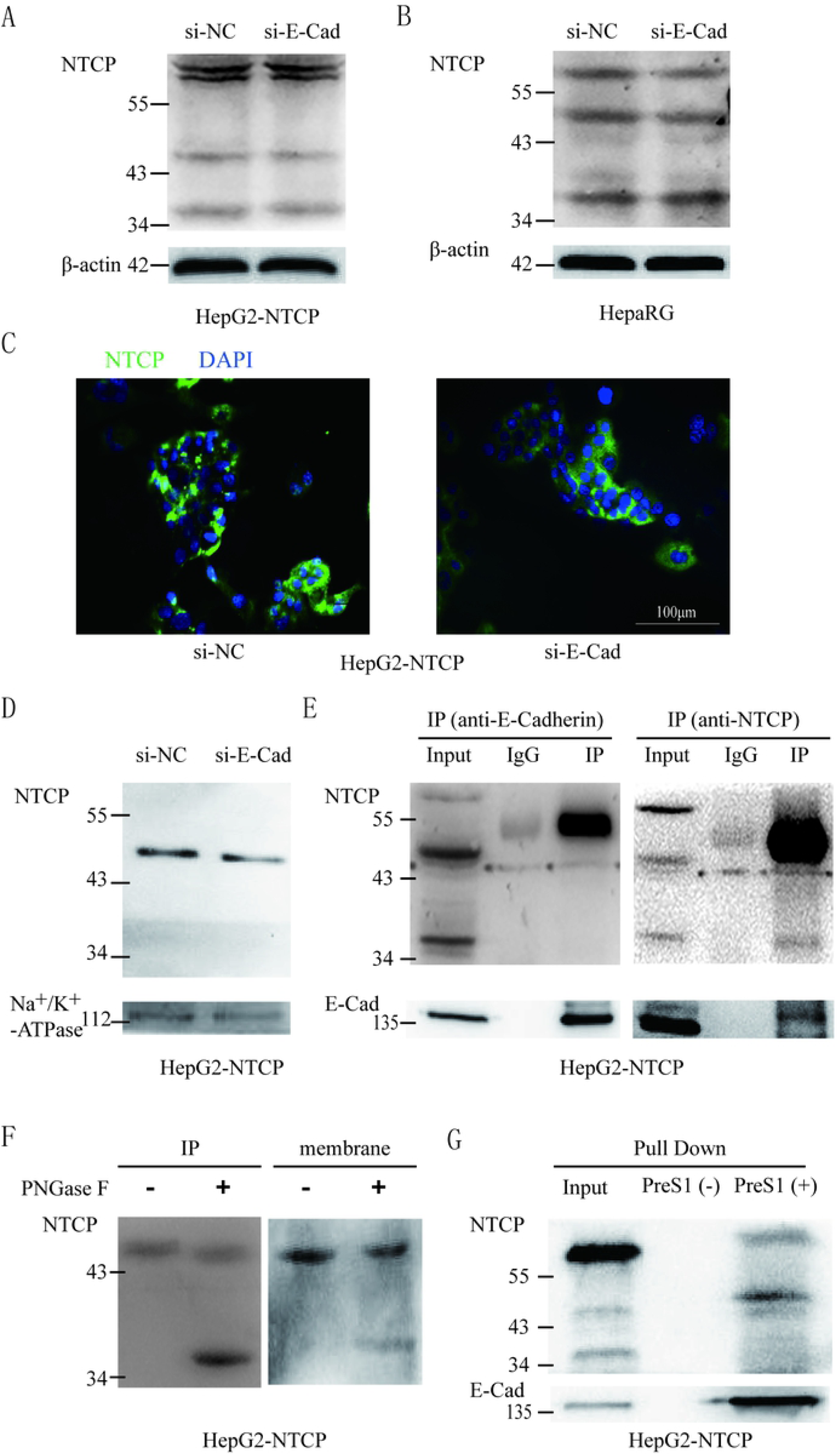
E-cadherin facilitates NTCP localization to the cell surface through interacting with glycosylated NTCP. Total protein was extracted three days post-transfection with siRNA-NC or siRNA-E-cadherin. NTCP expression was assessed by western blot analysis in (A) HepG2-NTCP and (B) HepaRG cells. (C) NTCP expression was assessed by immunofluorescence three days post-transfection with E-cadherin siRNA in HepG2-NTCP cells. (D) Membrane protein was extracted three days post-transfection with siRNA-NC (left panel) or siRNA-E-cadherin (right panel) and NTCP expression was assessed by western blot analysis in HepG2-NTCP cells. (E) Co-immunoprecipitation was performed to confirm the interaction between E-cadherin and NTCP. Input: Total protein from cell extract. IgG of rabbit was used as the control group. IP: Total protein from Hep2-NTCP cells was incubated with anti-E-cadherin, or anti-NTCP at 4 °C overnight. (F) The precipitate of IP and membrane protein were treated by PNGase F at 37 °C for 1 h and detected by western blot. (G) The HepG2-NTCP cell lysates were incubated with pre-S1 to confirm that preS1 was bound to glycosylated NTCP. Representative data is shown from triplicate experiments.

To elucidate the mechanism employed by E-cadherin to impact the cellular distribution of NTCP, we further examined the interactions between E-cadherin and NTCP via co-immunoprecipitation (co-IP). Surprisingly, when an E-cadherin antibody was used to precipitate the cellular lysate of HepG2-NTCP cells, an approximately 50 kDa protein band was detected in the precipitates with the NTCP antibody. To confirm whether the 50 kDa protein was glycosylated NTCP, the precipitate was treated by PNGase F and further probed with NTCP antibodies. Results demonstrated that PNGase F treatment caused the 50 kDa protein to change into a 39 kDa variant; suggesting the 50 kDa protein was a glycosylated form of NTCP (Fig 7F). Furthermore, after treating cell membrane proteins with PNGase F, we also observed a shift in electrophoresis size from 50 kDa to 39 kDa, indicating that a large fraction of the NTCP localized at the cell membrane was glycosylated (Fig 7F). Lastly, after incubating preS1 with HepG2-NTCP cell lysates, we observed the formation of preS1, glycosylated NTCP and E-cadherin complexes (Fig 7G). Taken together these results suggest that E-cadherin binds to glycosylated NTCP, allowing for efficient localization to the cell surface.

### E-cadherin modulates hepatocyte polarization

Cell polarity has also been reported to impact the efficiency of viral infections [14]. As an intercellular adhesive molecule, E-cadherin has a central role in maintaining normal cell structure and, thus, may also affect hepatocyte polarization. To examine the relationship between E-cadherin and hepatocyte polarization, we determined via immunofluorescence that silencing of E-cadherin downregulated the expression of ZO-1, which is a tight junction marker, and regulated the distribution of multidrug resistance protein 2 (MRP2), another polarization marker (Fig 8A and 8B). Furthermore, western blot analysis confirmed that silencing of E-cadherin significantly downregulated the expression of ZO-1, however, had no apparent impact on MRP2 expression (Fig 8C). These results suggest that E-cadherin modulates hepatocyte polarization.

**Fig 8.**
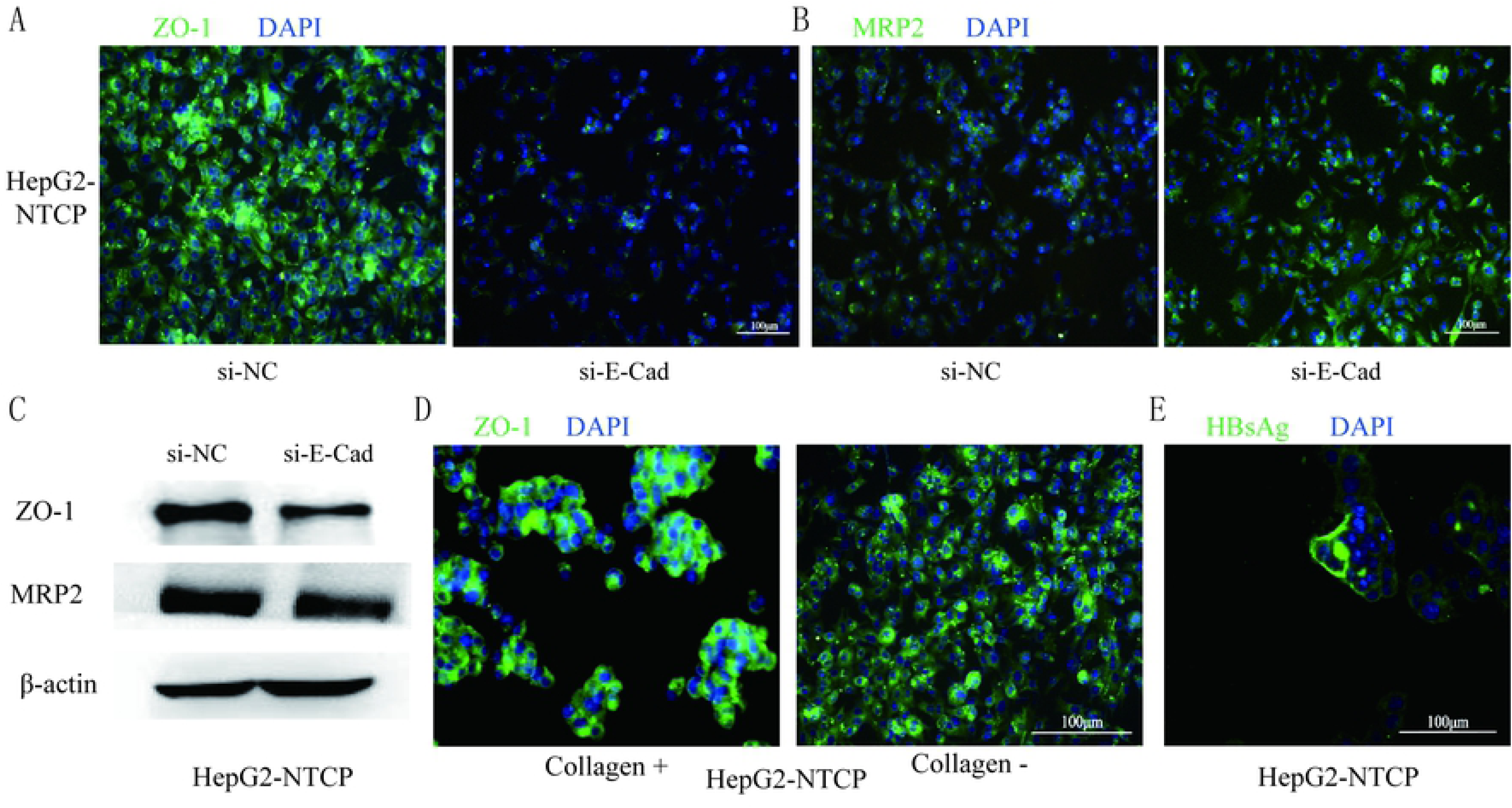
E-cadherin modulates hepatocyte polarization. The distribution and expression of (A) ZO-1 and (B) MRP2 were assessed by immunofluorescence and (C) western blot analysis three days post-transfection with siRNA-NC or siRNA-E-cadherin in HepG2-NTCP cells. (D) HepG2-NTCP cells were cultured in collagen-coated or normal culture dishes. The growth distribution of ZO-1 was assessed by immunofluorescence for different patterns including group and dispersion. Cells that were cluster grown in normal plates exhibited dispersed distribution of ZO-1 at the center of the hepatic island like that of in cells that were grown in collagen-coated plates and linear distribution at the edge of the hepatic island. (E) HepG2-NTCP cells in cluster growth were infected with HBV particles enriched from supernatant of HepaAD38 and the expression of HBsAg was assessed by immunofluorescence three days after infection. Representative data is shown from triplicate experiments.

HepaRG cells are only susceptible to HBV infection once they have differentiated into hepatic and biliary-like cells following treatment with DMSO [15]. Andreas Schulze et al. reported that hepatocyte-like HepaRG cells exhibited two unique patterns of MRP2 distribution namely, canalicular and dispersed. Moreover, elevated HBV infection rates were observed in hepatocyte-like cells exhibiting canalicular distribution patters of MRP2 [16]. We, therefore, also wanted to determine whether different culture conditions affected polarization and infection efficiency of HepG2-NTCP cells in different ways. Cells that were grown in collagen-coated plates exhibited diffuse distribution of ZO-1. Alternatively, cells that were cluster grown in normal plates exhibited dispersed distribution of ZO-1 at the center of the hepatic island and linear distribution at the edge of the hepatic island (Fig 8D). These results revealed that cells at the edge of the hepatic island were more susceptible to polarization. Further, we also determined that following infection of HepG2-NTCP cells with enriched HBV particles, cells at the edge of the hepatic island were more susceptible infection (Fig 8E).

## Discussion

HBV entry into the host cell is the critical step in their life cycle. This process includes particle delivery and capture, complex internalization, and membrane fusion [17]. HBV entry requires a tightly coordinated group of specific viral proteins and multiple host receptors [18]. In the current study, we report that HBV-expression upregulates E-cadherin expression, which acts as a novel host factor that facilitates HBV entry. The functional role of E-cadherin as a host entry factor was confirmed by numerous studies. Specifically we showed that silencing of E-cadherin acted to inhibit HBV particle entry into HepG2-NTCP, HepaRG and PHH cells (Fig 2 and 3); overexpression of E-cadherin contributed to HBV particle entry in HepaRG cells (Fig 4) and silencing of E-cadherin inhibited HBVpps entry in HepG2-NTCP and HepaRG cells (Fig 5). Moreover, E-cadherin silencing caused a significant decrease in the binding and internalization of the HBV pre-S1 peptide by HepG2-NTCP and PHH cells (Fig. 6). Mechanistic studies suggested that E-cadherin regulates the cell-surface distribution of NTCP (Fig 7) and affects hepatocyte polarization (Fig 8). This study represents an important step forward in understanding the molecular mechanisms and cellular regulatory events involved in HBV entry.

Previous studies reported that HBx or HBV inhibit E-cadherin expression. However, contrary to these reports, we have shown that the expression of E-cadherin was upregulated in cells that transiently or stably expressed HBV (Fig 1). Additionally, Wang et al. also reported that E-cadherin expression was downregulated following inhibition of HBV replication via silenced by siRNAs in HepG2.2.15 cells [19]. Similar contradictory results have been reported in HCV infection cell models [20]. However, the reason for these controversial results remains unclear and, thus, is it necessary to examine the expression level of E-cadherin in HBV-infected liver tissues that are free from carcinomas, to clearly elucidate the effect of HBV on E-cadherin expression.

Although Li et al. reported that HepG2-NTCP cells exhibited poorly susceptibility to HBV particles derived from patient serum-derived [21], our study demonstrated the opposite effect with HepG2-NTCP cells being efficiently infected with HBV particles from serum of chronic HBV patients. These results were confirmed by quantifying specific markers of HBV infection (HBV 3.5 kb RNA, HBc and HBsAg) to confirm infection of HepG2-NTCP cells (Fig 3). Similarly, Yan et al. successfully infected HepG2-NTCP cells with an HBV genotype B virus from the plasma of an HBV chronic carrier [6]. An additional study also reported successful infection of HepG2-NTCP cells with an HBV genotype D virus isolated from serum to determine the effect of glypican-5 (GPC5) expression on HBV entry [11].

E-cadherin is a type I classical cadherin and a calcium-dependent adhesion glycoprotein expressed in the epithelium [22]. E-cadherin has been described as a critical component in the regulation of pathways highly associated with cancer development, including cellular proliferation, apoptosis, invasiveness, metabolism, and metastasis, through mediating multiple cellular signaling pathways [23]. Many additional studies have focused on the effect that E-cadherin has on pathogenic infections caused by bacteria and viruses [24]. *Listeria monocytogenes* is a foodborne pathogen that crosses the intestinal barrier upon binding between its surface protein InlA and the host receptor E-cadherin occurs [25]. *Candida albicans* produces two types of invasins, namely, Als1 and Als3, in combination with E-cadherin or N-cadherin, which together facilitate the internalization of bacteria [26]. Moreover, E-cadherin associates directly with nectin-1 which is critical for HSV-1 infection [27]. Connolly et al. also reported that E-cadherin overexpression may strengthen intercellular junctions and shorten the distance between infected and uninfected cells, thereby increasing the regional concentration of nectin-1 to facilitate efficient viral spread [28]. E-cadherin was also found to play a critical role in HCV entry through regulating membrane distribution of the HCV receptors, claudin-1 (CLDN1) and occludin (OCLN) [20]. These studies provide evidence that E-cadherin acts directly as a receptor while also modulating the expression and activity of other receptors to mediate pathogenic infections.

Herein we show that E-cadherin plays a critical role in HBV entry through influencing the distribution of NTCP, the functional receptor of HBV. NTCP, a 349 residue glycoprotein, becomes glycosylated at N5 and N11 in the endoplasmic reticulum and Golgi apparatus before being trafficked to the cell surface and the band is from 39 kDa to 56 kDa [29]. Surprisingly, we have also discovered that E-cadherin interacts exclusively with the glycosylated form of NTCP (Fig 7E). A previous study indicated that glycosylation is essential for NTCP to act as a receptor for HBV since the non-glycosylated form of NTCP is rapidly internalized and degraded [29]. Yan et al. also found that Myr-preS1 cross-links with glycosylated NTCP [6]. However, another study claimed that both the glycosylated and non-glycosylated forms of NTCP effectively mediate HBV infection, since differentiated HepaRG cells only express non-glycosylated NTCP [30]. In our study, both glycosylated and non-glycosylated NTCP were detected in differentiated HepaRG and HepG2-NTCP cells, however, only glycosylated NTCP were detected in membrane proteins. Moreover, Myr-preS1 was found to bind exclusively to the glycosylated form (Fig 7). Our results showed that E-cadherin was exclusively associated with glycosylated NTCP (Fig 7B), and, thus, we propose that E-cadherin exerts a regulatory role on the cellular entry of HBV through interacting with glycosylated NTCP and facilitating its membrane localization in hepatocytes.

Furthermore, E-cadherin is the primary component of adherens junctions at the basolateral surfaces of polarized epithelial cells and function to establish cellular polarity [31]. Madin-Darby Canine Kidney (MDCK) cells have been shown to completely lose polarity in the absence of calcium ions in culture medium. However, this effect was reversed following addition of calcium ions, indicating that E-cadherin is a critical factor for establishing and maintaining epithelial cell polarization [32]. Lu et al. found that silencing of E-cadherin expression reduces the stability of tight junctions thereby destroying hepatocyte polarization, by directly reducing the expression of PAR3 [33]. Alternatively, Théard D et al. reported that downregulation of E-cadherin and β-catenin on the cell membrane in HepG2 cells did not significantly affect the formation of tight junctions, allowing for normal establishment of normal bile duct-like apical and basement membrane polarization structures; suggesting that E-cadherin is not essential for establishing hepatocyte polarization [34]. In our current study, silencing of E-cadherin served to reduce the expression of ZO-1, a tight junction protein located in the lateral membrane, while also affecting the distribution pattern of MRP2, a marker that is selectively transported to the apical cell membrane (Fig 8A-C). ZO-1 is bound to αE-catenin, which forms a E-cadherin/β-catenin/αE-catenin complex, thus, downregulation of ZO-1 would cause loss of tight junction formation [35]. These results clearly indicate that E-cadherin affects cell polarization.

Cell polarization, defined as the asymmetric distribution of components and functions in a cell, is not only required for proper cellular functioning but has also been defined as being associated with pathogenic infection [14]. The archetypal polarized animal cell is the epithelial cell, however, hepatocytes are also highly polarized. Previous studies have shown that polarized epithelial cells use tight junctions to create physical and immunological barriers to prevent the invasion of pathogens [14]. Pathogens such as *Helicobacter pylori* [36], *Salmonella typhimurium* [37], *Shigella dysenteriae* [38], and Rotavirus [39], either interfere with the establishment of cellular polarization, or adapt to use polarized molecules as their functional receptors, thereby facilitating self-infection of host cells. Additionally, cell polarization has been shown to facilitate the membrane distribution of receptors for specific pathogens, thereby acting as an essential component in the infection process. Specifically, the Reovirus surface protein, σ1 of Reovirus binds to the N-terminus of JAM-A to assist virus entry [39]. Coxsackie virus enters cells following binding of CAR as a co-receptor [40]. HCV adheres to the cell surface by firstly binding to CD81 and occludin, and subsequent binding to claudin to allow for cell entry [41]. Furthermore, Schulze et al. reported that hepatocyte polarization is essential for HBV entry [18]. This group also determined that infected HepaRG cells within hepatic islands are predominantly located at the edges. In our study, we found that the distribution of partially polarized molecules differs according to the specific culture conditions. ZO-1, located at the edges of the HepG2-NTCP cluster were found to exhibit linear polarity on the cell membrane (Fig 8D). Importantly, these HepG2-NTCP cells at the edges of the cluster were found to be more susceptible to HBV infection (Fig. 8E). These results suggest that polarized hepatocytes are more susceptible to HBV infection, which is consistent with the results of previous studies.

In summary, HBV was found to upregulate the expression of E-cadherin, which bound glycosylated NTCP thereby, impacting the membrane distribution of NTCP and facilitating the establishment of hepatocyte polarity, serving to improve the rate of HBV entry. This study provides novel insights that advance the current understanding of the HBV life cycle as well as inform the development of pharmaceutical interventions targeting E-cadherin as a means to prevent HBV infection.

## Materials and Methods

### Cell Lines

HepG2, HepG2-NTCP, HepAD38 cell lines (in which HBV replication can be regulated by tetracycline), Huh-7 and HepG2.2.15, were provided by Professor Juan Chen of the Chongqing Medical University, Chongqing, China. HepG2 cells were cultured in Modified Eagle’s Medium (MEM, Hyclone, USA) with 10 % fetal bovine serum (FBS, Gibco, USA). HepG2-NTCP and HepAD38 cells were maintained in Dulbecco’s Modified Eagle Medium (DMEM, Hyclone, USA) supplemented with 10% FBS and 400 µg/mL of G418. HepG2-NTCP cells were cultured with or without collagen pretreatment. Huh-7 and HepG2.2.15 cells were maintained in DMEM containing 10 % FBS. HepaRG cells were purchased from Beijing Beina Science and Technology, China and were cultured in William’s E Medium (Sigma-Aldrich, USA) with 10 % FBS, 2 mM L-glutamine, 5 ug/mL insulin (Sigma-Aldrich, USA) and 50 μM hydrocortisone hemisuccinate. Primary human hepatocytes (PHH) were obtained from ScienCell Research Laboratories (Carlsbad, CA, USA) and were maintained in Hepatocyte Medium (HM; Catalog no. 5210; ScienCell Research Laboratories, Carlsbad, CA, USA). All cells were cultured in a humidified incubator at 37 °C with 5% CO_2_.

### Plasmids and gene products

The pNL4-3.Luc.R-E-reporter plasmid was obtained through the NIH AIDS Reagent Program. The pcDNA3.1-E-cadherin was obtained from addgene, while the pcDNA3.1-HBV1.1 (replication-competent 1.1-fold overlength HBV), and pcDNA3.1-HBV1.3 (replication-competent 1.3-fold overlength HBV) plasmids were provided by Professor Juan Chen of the Chongqing Medical University, Chongqing, China. To study the effect of E-cadherin on HBV binding to HepG2-NTCP cells, the FITC-labeled preS1 peptide encoding the stable region of preS1 (2−47aa) was synthesized by Zhejiang Baitai Biological Company, China. The sequence used to produce the FITC (or Biotin)-labeled preS1 peptide was: Myr-GTNLSVPNPLGFFPDHQLDPAFGANSNNPDWDFNPNKDHWEANQVK [6].

### Protein extraction and western blot analysis

Total protein content was extracted from cells using a Protein Extraction Kit (KaiJi, China) according to manufacturer’s instructions. Protein content was quantified via bicinchoninic protein assays (Bio-Rad, China) according to manufacturer’s instructions. Equal amounts of total protein from each sample (50 μg) were separated using sodium dodecyl sulfate-polyacrylamide gel electrophoresis (SDS-PAGE) and transferred to a nitrocellulose membrane. The membranes were incubated with polyclonal rabbit anti-HBc (1:1,000; Dako, USA), rabbit anti-NTCP (1:1,000; Sigma-Aldrich, USA), rabbit anti-ZO-1 (1:1,000; Proteintech, China) rabbit anti-MRP2 (1:1000; Cell Signaling Technology, USA), and anti-actin (1:1;000; Bio-Rad, China) primary antibodies at 4 °C overnight, and subsequently incubated with horseradish peroxidase (HRP)-conjugated secondary antibodies (1:10,000; Bio-Rad, China) at 37 °C for 1 h. Blots were developed using an enhanced chemiluminescence reagent (Pierce; USA) according to manufacturer’s instructions.

### Preparation of cell membrane fractions

Cell membrane proteins were isolated using MinuteTM Plasma Membrane Protein Isolation and Cell Fractionation Kit (ScienCell Research Laboratories, USA) according to manufacturer’s instructions. Protein concentrations in the extraction samples were quantified using BCA protein assays (Bio-Rad, China) according to the manufacturer’s instructions.

### Immunofluorescence assay

Coverslips containing cells were washed with phosphate buffered saline (PBS) twice, fixed with 4 % paraformaldehyde at 37 °C for 10 min, blocked with goat serum at 37 °C for 1 h, and incubated with primary antibodies at 4 °C overnight. The coverslips were then incubated with secondary antibodies (FITC-anti-rabbit IgG) at 37 °C for 1 h, and stained with 4′,6-diamidino-2-phenylindole (DAPI) at 37 °C for 8 min. Immunofluorescence analysis were performed using a Leica DFC5000 camera attached to an Olympus FV500 Confocal Laser Scanning microscope.

### Analysis of messenger RNA expression by quantitative real-time polymerase chain reaction

Total RNA was extracted from cells using Trizol reagent (Invitrogen, USA) according to the manufacturer’s instructions. Reverse-transcription of total RNA was performed using PrimeScript RT Reagent Kit with gDNA Eraser (Takara; Japan). Quantitative real-time polymerase chain reaction (qRT-PCR) of the HBV 3.5 kb mRNA was performed using SYBR Green Master Mix (Roche; Germany). The primers of the HBV 3.5 kb were, forward: 5’-GCCTTAGAGTCTCCTGAGCA-3’ and reverse: 5’-GAGGGAGTTCTTCTTCTAGG-3’. The primers of the β-actin were, forward: 5’-TCCCTGGAGAAGAGCTACGA-3’ and reverse: 5’-AGCACTGTGTTGGCGTACAG-3’. All values were normalized to β-actin expression and were calculated using the 2^ΔΔCT^ method.

### siRNA transfections

Targeting small interfering (siRNAs) or nontargeting siRNAs were transfected into HepG2-NTCP, HepaRG or PHH cell lines using siRNA-mate (Genephama, China) according to the manufacturer’s protocol. Further treatments were typically performed 72 h after siRNA transfection began, when gene silencing was determined to reach peak efficiency. The sequence used for siRNA targeting E-cadherin was: 5′-CAGACAAAGACCAGGACUA-3′. The sequence of the siRNA targeting NTCP was: 5′-CAGUUCUCUCUGCCAUCAA-3′. The sequence of the siRNA targeting HBV was: 5′-GCGGGUAUAUUAUAUAAGATT-3′. The sequence of the negative control siRNA was: 5′-UUCUCCGAACGUGUCACGUdTdT-3′.

### HBV particles production and infection

HBV particles were enriched from the supernatant of HepAD38 cells using 8 % PEG8000 (Sigma, USA) [42]. HepG2-NTCP, HepaRG and PHH cells were seeded in 24-well plates and maintained in William’s E Medium supplement with 10 % FBS, 2 mM L-glutamine, and 2 % dimethylsulfoxide (DMSO) for 1 d at the time of HBV infection. HBV infection was assessed at 2 d by qRT-PCR quantification of HBV 3.5 kb RNA, and at 3 d via western blot analysis of HBV core protein (1:1,000; Dako, USA), and by immunofluorescence assay of HBsAg, using the rabbit anti-HBsAg antibody (1:100; Abcam, USA).

HBV particles from adult chronic HBV patients were obtained and infected as previously described [9]. Briefly, HepG2-NTCP and PHH cells were seeded at a density of 5 × 10^4^ per well into 24-well plates. On the following day, cells were incubated with the HBV-positive serum (>5 × 10^7^ copies/mL) diluted in DMEM at a MOI of 10. Infected cells were maintained in DMEM or HM with 2 % FBS. The HBV 3.5kb RNA, HBc and HBsAg were assessed four days post infection.

### Packaging and infection of HBV pseudotyped particles

HBV pseudotyped particles (HBVpps) could infect liver cells but cannot replicate in human hepatocytes. Infection and packaging of HBVpps were achieved as previously described [43]. Briefly, HBVpps were produced by co-transfecting pNL4-3.Luc.R-E- and pcDNA3.1-HBV1.3 plasmids into Huh7 cells using Lipofectamine 2000 Transfection Reagent (Invitrogen, USA). The supernatants containing HBVpps were harvested 72 h after transfection and centrifuged at 1,500 *× g* for 15 min to remove debris. The pseudotyped particles were then added to 96-well microplates, which had been seeded with target cells at a density of 5 × 10^3^ on the previous day, at a volume of 100 µL/well. The media was supplemented with polybrene (Santa Cruz, USA) at a final concentration of 4 µg/mL. Twenty-four hours after infection, 100 μL of fresh media was changed per well. Luciferase assays were performed after 72 h using a GloMaxTM 96 Microplate Luminometer (Promega, USA).

### Co-immunoprecipitation assay

Cells were lysed in immunoprecipitation buffer supplemented with complete protease inhibitor. After lysing with ultrasound, lysates were centrifuged to remove insoluble components. Lysates were then incubated overnight with rabbit anti-E-cadherin (Cell Signaling Technology, USA) or rabbit IgG primary antibodies at 4 °C. Protein G beads (Beyotime, China) were then added to the lysates. The precipitated proteins were measured via western blot analysis as described above.

### Pull-down Assay

Magnetic beads (Thermo Scientific) were incubated with biotin-labeled preS1 antibodies at 37 °C for 1 h according to manufacturer’s instructions. The beads were washed thrice with TBST, incubated with 500 uL HepG2-NTCP cell lysate at 4 °C overnight, and washed a further three times. Thirty microliters of SDS-PAGE reducing sample buffer was then added to each tube and heated at 100 °C in a heating block for 5 min. The precipitated proteins were quantified via western blot analysis.

### Ethics Statement

We collected fresh serum from adult chronic HBV patients. Written informed consent was obtained from each subject. The study was accepted by the Ethics Committee of the Second Affiliated Hospital of Chongqing Medical University.

### Statistical analyses

Statistical Package for the Social Sciences (SPSS) 16.0 was used to perform unpaired Student’s t-tests. *p* values < 0.05 were considered statistically significant. Each experiment was repeated a minimum of three times.

## Acknowledgments

The following reagent was obtained through the NIH AIDS Reagent Program, Division of AIDS, NIAID, NIH: pNL4-3.Luc.R–.E– from Dr. Nathaniel Landau.

